# Dynamic properties in functional connectivity changes and striatal dopamine deficiency in Parkinson’s disease

**DOI:** 10.1101/2024.03.06.583342

**Authors:** Adrian L. Asendorf, Hendrik Theis, Marc Tittgemeyer, Lars Timmermann, Gereon R. Fink, Alexander Drzezga, Carsten Eggers, Merle C. Hoenig, Thilo van Eimeren

## Abstract

**Introduction:** Recent studies in Parkinson’s disease (PD) patients reported disruptions in dynamic functional connectivity (dFC, i.e., a characterization of spontaneous fluctuations in functional connectivity over time). Here, we assessed whether the integrity of striatal dopamine terminals directly modulates dFC metrics in separate PD cohorts, indexing dopamine-dependent changes in large-scale brain network dynamics and its implications in clinical features.

**Methods:** We pooled data from two cohorts reflecting early PD. From the Parkinson’s Progression Marker Initiative (PPMI) cohort, resting-state functional magnetic resonance imaging (rsfMRI) and dopamine transporter (DaT) SPECT were available for 63 PD patients and 16 age- and sex-matched healthy controls. From the clinical research group 219 (KFO) cohort, rsfMRI imaging was available for 52 PD patients and 17 age- and sex-matched healthy controls. A subset of 41 PD patients and 13 healthy control subjects additionally underwent ^18^F-DOPA-PET imaging. The striatal synthesis capacity of ^18^F-DOPA PET and dopamine terminal quantity of DaT SPECT images were extracted for the putamen and the caudate. After rsfMRI pre-processing, an independent component analysis was performed on both cohorts simultaneously. Based on the derived components, an individual sliding window approach (44s window) and a subsequent k-means clustering were conducted separately for each cohort to derive dFC states (reemerging intra- and interindividual connectivity patterns). From these states we derived temporal metrics, such as average dwell time per state, state attendance, and number of transitions and compared them between groups and cohorts. Further, we correlated these with the respective measures for local dopaminergic impairment and clinical severity.

**Results:** In both cohorts, dFC analysis resulted in three distinct states, varying in connectivity patterns and strength. In the PPMI cohort, PD patients showed a lower state attendance for the globally integrated (GI) state (X^2^(1, N=79) = 5.82, *p*= 0.016) and a lower number of transitions (U(N=79) = 337.5, z = −2.06 p= .039) than controls. Significantly, worse motor scores (UPDRS-III) and dopaminergic impairment in the putamen and the caudate were associated with low average dwell time in the GI state (UPDRS-III: τ_b_(N=63) = −.281; *p* =.003, DaT putamen: τ_b_(N=63)=.213, *p*= .023, DaT caudate: τ_b_(N=63)=.209, *p*= .025) and a low total number of transitions (UPDRS-III: τb(N=63)= −.308; p = .001, DaT putamen: τ_b_(N=63)=.350, *p*<.001, DaT caudate: τ_b_(N=63)=.251, *p*=.007). Additionally, worse motor performance was associated with a low number of bi-directional transitions between the GI and the lesser connected (LC) state (τ_b_(N=63)= −.237; *p =*.019). These results could not be reproduced in the KFO cohort: No group differences in dFC measures or associations between dFC variables and dopamine synthesis capacity or clinical measure were observed.

**Conclusion:** In early PD, relative preservation of motor performance may be linked to a more dynamic engagement of an interconnected brain state. Specifically, those large-scale network dynamics seem to depend on striatal dopamine availability. Notably, we obtained these results in only one cohort, but not in a replication sample.

**Key points:**

- Exploring dopamine’s role in brain network dynamics in two Parkinson’s disease (PD) cohorts, we unraveled PD-specific changes in dynamic functional connectivity (dFC).
- In the discovery cohort, results suggest striatal dopamine availability influences large-scale network dynamics that are relevant in motor control.
- In the confirmation cohort, these findings were not replicated, indicating PD-specific dFC changes are dependent on unrecognized cohort features.

## Introduction

Parkinson’s disease (PD) is characterized by a degeneration of the nigrostriatal dopaminergic neurons, causing a wide range of motor and non-motor symptoms. Region-specific associations have revealed a strong relationship between striatal dopamine availability, motor symptom severity (Asenbaum et al., 1997; Benamer et al., 2000; Pirker, 2003), and higher-order cognitive tasks (Cools, 2011; Costa et al., 2014). From a network-based perspective, this suggests dopamine is a crucial player in effectively recruiting large-scale networks associated with these cognitive and motor domains. Using resting-state functional MRI (rsfMRI) to investigate connectivity across different networks has repeatedly shown that the putamen, which is known to express the most prominent striatal dopamine reduction in clinical PD, loses connectivity strength with frontal brain structures (Rolinski et al., 2016). Further, in recent multimodal approaches incorporating dopamine positron emission tomography (PET) or dopamine transporter (DaT) single-photon emission computed tomography (SPECT), it has been illustrated that the degeneration of nigrostriatal connectivity functionally impairs distinct striatocortical (Ruppert et al., 2020) and putamen-midbrain (Rieckmann et al., 2015) projections.

Additionally, an increase in dopaminergic deficit was associated with decreased putaminal interconnectivity over time (Li et al., 2020). These studies suggest that dopamine is fundamental for the efficient interplay between motor processing areas and parts of the basal ganglia. Interestingly, PD-specific changes in large-scale network organization normalize after taking dopaminergic medication (P. T. Bell et al., 2015; Cole et al., 2013; Wu et al., 2009). Thus, network consequences of dopamine loss may not be limited to brain regions directly connected to the basal ganglia. However, how striatal dopamine deficiency specifically perturbs the interplay of large-scale brain networks remains unresolved.

Notably, most of these studies assessing network organization modulated by dopamine capacity in PD have typically assumed temporal stability and, hence, time-averaged functional networks across the resting state data acquisition. However, functional networks dynamically fluctuate within scales of seconds and minutes (Chang & Glover, 2010; Lurie et al., 2020). Information contained within the temporal features of spontaneous functional connectivity fluctuations (dynamic functional connectivity, dFC) have been suggested to index changes in macroscopic neural activity patterns underlying critical aspects of cognition and behavior (Calhoun et al., 2014; Hutchison et al., 2013). In fact, compared to the static approach, growing evidence suggests such dFC metrics to be a sensitive indicator for determining disability level (Tozlu et al., 2021) and distinguishing between individuals with pathological conditions and those without (Jin et al., 2017; Rashid et al., 2016). Concomitantly, altered network dynamics have been reported for various neuro-psychiatric conditions (Damaraju et al., 2014; de Lacy et al., 2017; Jones et al., 2012; Kaiser et al., 2015; Liu et al., 2017; Sakoğlu et al., 2010).

In PD, several recent whole-brain dFC accounts suggested the relevance of network dynamics in the clinical presentation of PD (Cao et al., 2023; Cordes et al., 2018; Díez-Cirarda et al., 2018; Fiorenzato et al., 2019; Kim et al., 2017). Importantly, these studies could show that PD-specific changes in dFC were linked to motor symptom severity (Kim et al., 2017), mild cognitive impairment (MCI) (Díez-Cirarda et al., 2018; Fiorenzato et al., 2019), PD dementia (PDD) (Fiorenzato et al., 2019), and autonomic dysfunction (Cao et al., 2023). Together, these studies emphasize that motor and cognitive impairments in PD appear to be driven by spatial *and* temporal alterations of large-scale network dynamics. However, reports from PD are inconsistent. While some studies suggested PD to be associated with a higher occurrence of highly interconnected states (i.e., reoccurring patterns of functional connectivity across time) (Díez-Cirarda et al., 2018; Kim et al., 2017), others demonstrated PD (Fiorenzato et al., 2019) and PD progression (Cao et al., 2023) to be associated with a higher occurrence of sparsely connected states. These inconsistent findings, however, may be due to methodological differences in data analyses and study design. Moreover, the currently available studies employing dFC in PD are limited to correlations with behavior or clinical severity and do not use quantified information about the degree of dopaminergic terminal loss. Direct investigations of the modulatory role of dopamine on dynamic network fluctuations are currently missing.

To this end, we analyzed datasets of two age- and sex-matched cohorts. The Parkinson’s Progression Marker Initiative (PPMI) data set (https://www.ppmi-info.org/study-design/study-cohorts#overview/) included rsfMRI for 63 PD patients and 16 controls and DaT-SPECT as a measure for presynaptic dopaminergic dysfunction. In comparison, the KFO dataset (https://gepris.dfg.de/gepris/projekt/101434521?language=en) comprised 52 PD patients and 17 controls, including ^18^F-Dopa PET as the measure of presynaptic dopaminergic impairment. Notably, both PD groups consisted of individuals who either discontinued dopaminergic medication before the scan or were under no dopaminergic medication at all. The objectives of our study were as follows: First, to probe the reproducibility of our dFC analysis in two different PD cohorts. Second, to investigate the specific influence of dopamine deficiency on large-scale network dynamics. To achieve this, we obtained PD-specific dFC changes by comparing the dFC variables of a healthy control group with a PD group for each cohort separately. Consequently, we tested whether these PD-specific dFC changes were associated with clinical cognition, motor scores, and dopamine availability in the striatum. We suspected PD-specific changes in dFC to appear in both groups. According to the literature discussed above, we expected these to be associated with either motor or cognitive function and dopaminergic deficit in the striatum.

## Materials and Methods

### Participants

Data used for this study included two different cohorts. The first cohort included subjects enrolled in the Clinical Research Group 219 (KFO219) in Cologne. For the KFO cohort, rsfMRI and structural MRI data of 52 PD patients and 17 healthy controls were available. A subset of 41 PD patients and 13 healthy control subjects additionally underwent ^18^F-DOPA-PET imaging. The average time difference between MRI and ^18^F-DOPA-PET imaging was 29 days (SEM 8.05 days). All PD patients fulfilled the UK Brain Bank Criteria for PD (Gibb & Lees, 1988). All participants gave informed consent. The study was approved by the local ethics committee (EK12-265).

The second cohort was matched to the KFO cohort regarding age and sex and was composed of individuals registered in PPMI. The PPMI dataset included 63 PD patients and 16 age- and sex-matched healthy controls. These subjects 1) had undergone a fixed rsfMRI protocol (see MRI acquisition), 2) were between fifty and eighty-five years old, and 3) had data of structural MRI, UPDRS-OFF, and DaT-SPECT imaging available. In PD individuals, these data were not dated further from the rsfMRI as +- 182 days. The PD group’s average time difference between the DaT scan and rsfMRI acquisition was 12 days (SEM 2.29). For the control group, the difference was 728 days (SEM 165.22). All PD patients had a diagnosis of Parkinson’s disease and exhibited at least two of the following: resting tremor, bradykinesia, and rigidity.

Exclusion criteria for both cohorts included a history of other severe or neurological diseases and/or current medication that affects brain function, a clinical diagnosis of dementia, and MRI safety exclusion criteria. Any quality defects of the rsfMRI, like artifacts, phantoms, tilted planes, or cuts in the cortex, were excluded. The demographic characteristics of the two cohorts are summarized in Table 1.

**Table 1:**
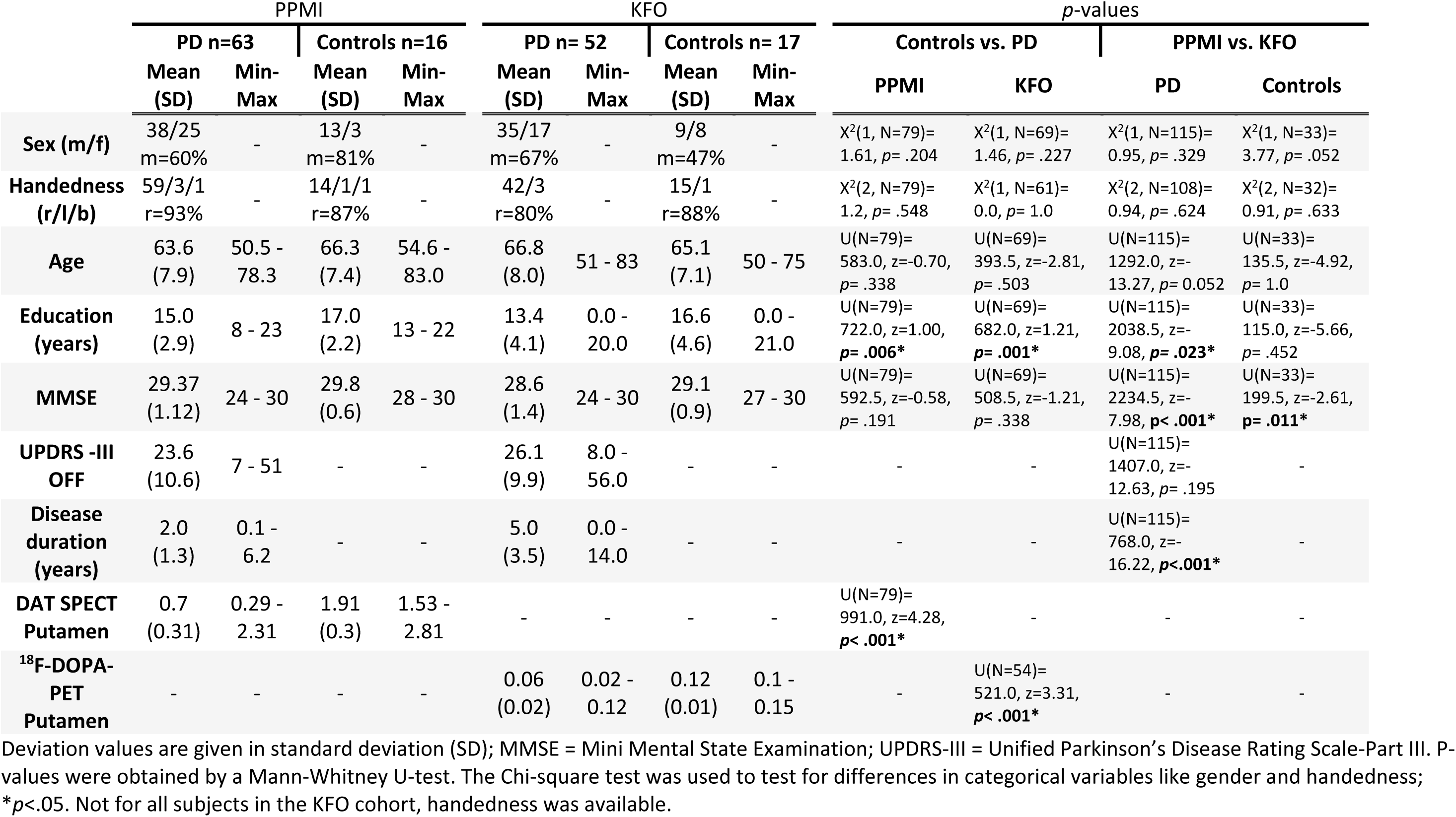
Demographic and clinical Data from both cohorts.

### Clinical and neuropsychological assessment

In both cohorts, disease severity in PD patients was assessed using the Unified Parkinson’s Disease Rating Scale (UPDRS) Part III (Goetz, 2003) in an unmedicated state (OFF). For the KFO dataset, this represented at least 12 hours of withdrawal of dopamine replacement therapy or 72 hours for extended release of dopamine agonists. In the PPMI cohort, 31 PD patients were taking medication. These subjects discontinued dopaminergic medication at least 6 hours (16 hours on average) before clinical examination (UPDRS-III). Furthermore, using the Mini Mental State Examination (MMSE) in the KFO dataset and the Montreal Cognitive Assessment (MoCA) in the PPMI cohort, cognitive function was tested. To establish a comparability of these two tests, we converted the MoCA scores to MMSE scores according to a recently introduced table (Fasnacht et al., 2023).

### Acquisition of the dopamine synthesis capacity

Measures of presynaptic dopamine deficiency were used to assess the possible impact of striatal dopamine availability on large-scale network connectivity dynamics. In the KFO cohort, ^18^F-DOPA-PET imaging was used, which was acquired for a subset of 56 subjects on a 24-detector ring scanner (ECAT EXACT HRRT, Siemens) at the Max-Planck-Institute for Metabolism Research in Cologne. ^18^F-DOPA-PET was performed in the morning after overnight fasting and OFF dopaminergic medication. The image processing procedure was previously described elsewhere (Hammes et al., 2019). Briefly, dopamine metabolism was assessed using a reference tissue model (Patlak plot) on dynamic scans comprising nine movement-corrected and spatially normalized frames. This data processing resulted in images exclusively representing the striatum’s presynaptic dopamine synthesis capacity. Finally, the mean presynaptic dopamine synthesis capacity was extracted from four striatal volumes of interest (VOIs): the left and right putamen and caudate nucleus.

In the PPMI cohort, DaT-SPECT imaging was carried out in different imaging centers on different scanners. DaT-SPECT imaging was performed according to the PPMI standardized protocol to quantify dopamine transporters in the striatum. Each imaging center reconstructed SPECT images using a standard iterative reconstruction algorithm and then corrected them for attenuation. Detailed imaging protocols can be comprehended here: https://www.ppmi-info.org/study-design/research-documents-and-sops

Our analysis used mean putamen and caudate values for both cohorts.

### MRI acquisition

An anatomical T1-weighted MRI and a rsfMRI series were acquired for each participant. In KFO, the structural T1-weighted images were obtained on a 3.0 T Siemens Magnetom Prisma scanner using the following acquisition parameters: repetition time (TR) = 2300 ms, echo time = 2.32 ms, flip angle = 8°, field of view =230 mm, slice thickness =0.90 mm, voxel size= 0.9 x 0.9 x 0.9 mm, number of slices = 192. The rsfMRI series included a gradient echo-planar imaging sequence with interleaved acquisition mode using the following parameters: TR = 0.776 ms, echo time = 37.4 ms, flip angle = 55°, field view =208 mm, voxel size = 2.0 x 2.0 x 2.0 mm, slice thickness = 2 mm. The acquisition time was 8 minutes and contained 617 time points comprised of 72 axial slices each.

In PPMI, the structural images were acquired on 3.0 T Siemens Magnetom scanners (Trio-A- Tim, Verio, or Prisma) using a slice thickness of 1 mm. The rsfMRI series used in the PPMI cohort was carried out in interleaved acquisition mode running the following protocol: TR = 2400 ms, echo time = 25.0 ms, voxel size = 3.3 x 3.3 x 3.3 mm, slice thickness 3.3 mm. The acquisition time was 8 minutes and 24 seconds, containing 210 time points composed of 40 axial slices each.

In both cohorts, subjects were asked to keep their eyes open and remain still.

### Functional MRI data preprocessing and motion control

Preprocessing was carried out using the CONN toolbox v20.b (Whitfield-Gabrieli & Nieto-Castanon, 2012) implemented in Matlab (Matlab R2020b Update 4, MathWorks, Inc., Natick, Massachusetts, United States). A default preprocessing pipeline was used to normalize the images to the MNI (Montreal Neurological Institute) space (web.conn-toolbox.org/fmri-methods/preprocessing-pipeline). The pipeline includes spatial realignment, slice-timing correction, outlier identification, segmentation, normalization, and smoothing (Gaussian kernel 8mm full-width half maximum).

Using the artifact removal tool included in CONN potential outlier scans caused by subject motion were identified, yielding six-dimensional motion vectors composed of translational (x, y, z axes) and rotational (pitch, yaw, roll) displacement scores. These six displacement scores were each transformed into single framewise displacement (FD) values using a published formula (Power et al., 2012). Acquisitions were discarded if they showed a mean FD > ½ voxel size (1 mm in KFO and 1.65 mm in PPMI) and/or a maximum displacement of more than one voxel (2 mm in KFO and 3.3 mm in PPMI). According to these criteria, four PD patients and two healthy controls were excluded from the KFO cohort, resulting in a global mean FD of 0.19 mm for the final cohort. Finally, two PD patients and two healthy controls were excluded from the PPMI cohort, which resulted in an average FD of 0.34 mm.

### Identification of intrinsic connectivity networks

#### Dimensionality reduction and group component definition

Independent group components were identified in a data-driven approach by means of data reduction and a spatial independent component analysis (ICA) utilizing the Group ICA fMRI toolbox (GIFT) (v3.0c, https://trendscenter.org/software/gift/). To obtain comparable components between the two cohorts, the group component identification was conducted in a pooled approach on all subjects of the two cohorts, involving healthy controls and PD patients (see Fig 2 left). The applied data reduction and ICA protocol followed previous dFC analyses (Allen et al., 2014; Damaraju et al., 2014; Díez-Cirarda et al., 2018; Fiorenzato et al., 2019; Kim et al., 2017): On the subject level, a principal components analysis (PCA) reduced the data to 120 principal components (PCs). At the group level, the concatenated reduced data was condensed to 100 PCs using the expectation maximization algorithm for PCA (Roweis, 1998). Following the data reduction step, a Infomax ICA algorithm (Himberg et al., 2004) decomposed the group-level data into independent networks (A. J. Bell & Sejnowski, 1995), producing a single set of 100 group independent components (ICs). Reliability through process stability was ensured by repeating the Infomax algorithm 20 times using the ICASSO method. Only components with a stability index > 0.9 were selected, leaving 76 ICs for spatial characterization. Thereafter, a GICA-based back-reconstruction step was added to compute individual subject-specific spatial maps required for further analysis (Calhoun et al., 2001).

#### Feature identification and thresholding

Of the 76 stable ICs, we identified 57 independent network components (INCs) utilizing identification criteria previously described (Allen et al. 2014, 2011): 1) peak activations located primarily in the grey matter, 2) low spatial overlap with known vascular and ventricular spaces, 3) low susceptibility for artifacts. The resulting 57 INCs were subsequently grouped into 14 previously established resting state networks (RSNs) (Shirer et al. 2012; Allen et al. 2014): Using the DICE similarity coefficient (a measure of spatial overlap), the INCs were sorted according to their highest spatial overlap with one of the RSNs. In seven instances, a single INC had a similar spatial overlap with several RSNs. For these cases, one of the RSN options was chosen by the shared decision of two experts (M.H., T.v.E.). An overview of the component patterns and corresponding RSNs can be found in Figure 1. With the preceding steps, including data reduction, ICA, feature identification, and back-reconstruction, we defined RSNs that share comparable activation patterns and anatomical locations across each individual and both cohorts. This approach enabled us to assess changes in RSN connectivity and compare these changes between individuals, groups, and cohorts.

**Figure 1:**
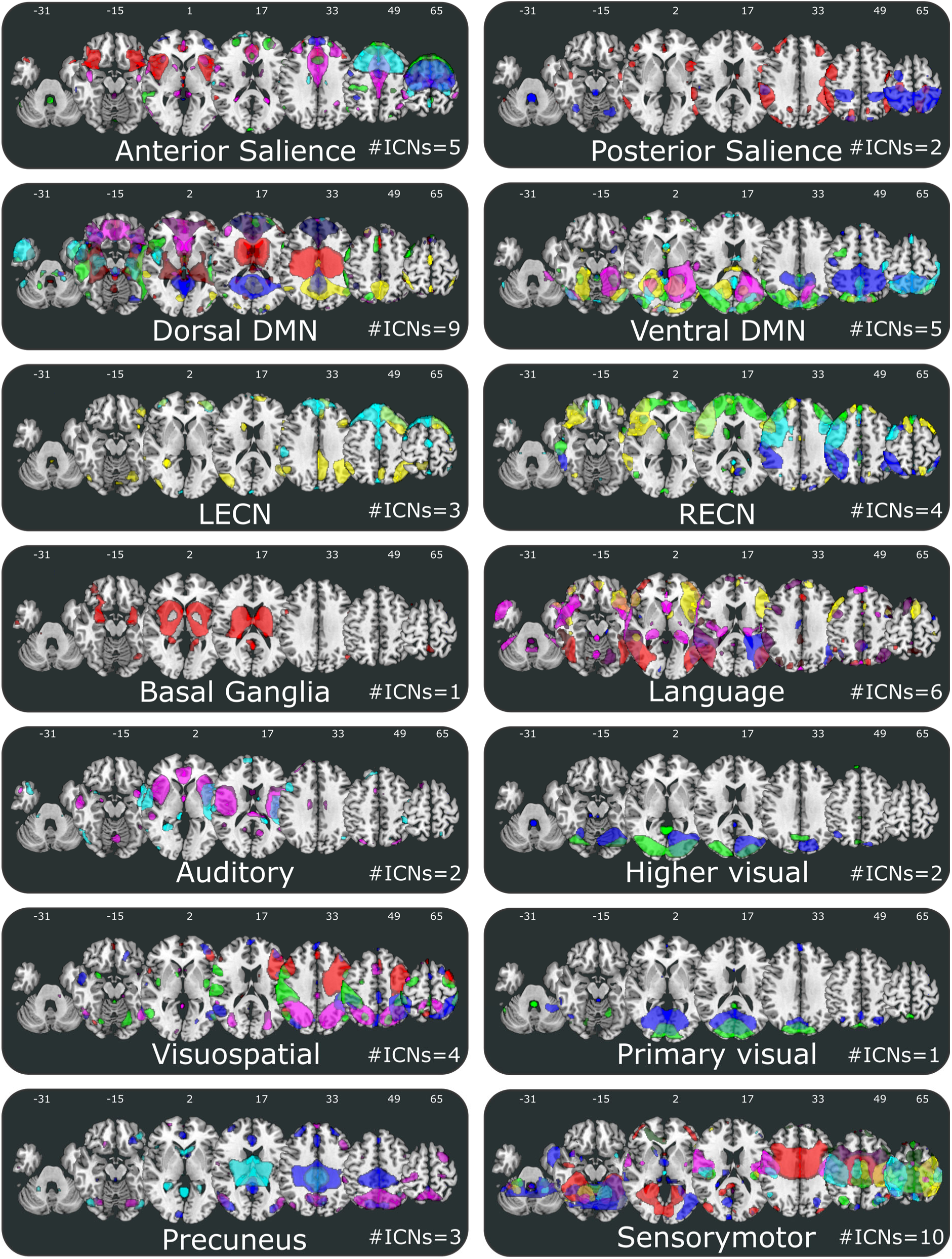
Composite maps of the 57 independent network components (INCs) identified by group ICA. Based on their anatomical and functional properties, 57 INCs were grouped into the 14 resting-state networks (RSNs) for functional connectivity previously established by Shirer et al., 2012. Each color in the composite maps corresponds to a single INC. The number of ICNs sorted into each RSN is stated in the bottom right corner of each panel. DMN = default mode network; LECN = left executive network; RECN = right executive network.

### Dynamic functional connectivity analysis

The dynamic functional connectivity (dFC) analysis was conducted separately for each cohort. Using the temporal dFNC toolbox in GIFT, we combined two approaches: The sliding window approach, which determines changes in functional connectivity across time in single subjects, and the *k*-means clustering algorithm, which enables the extraction of reoccurring patterns of functional connectivity across time (see Figure 2 right).

**Figure 2:**
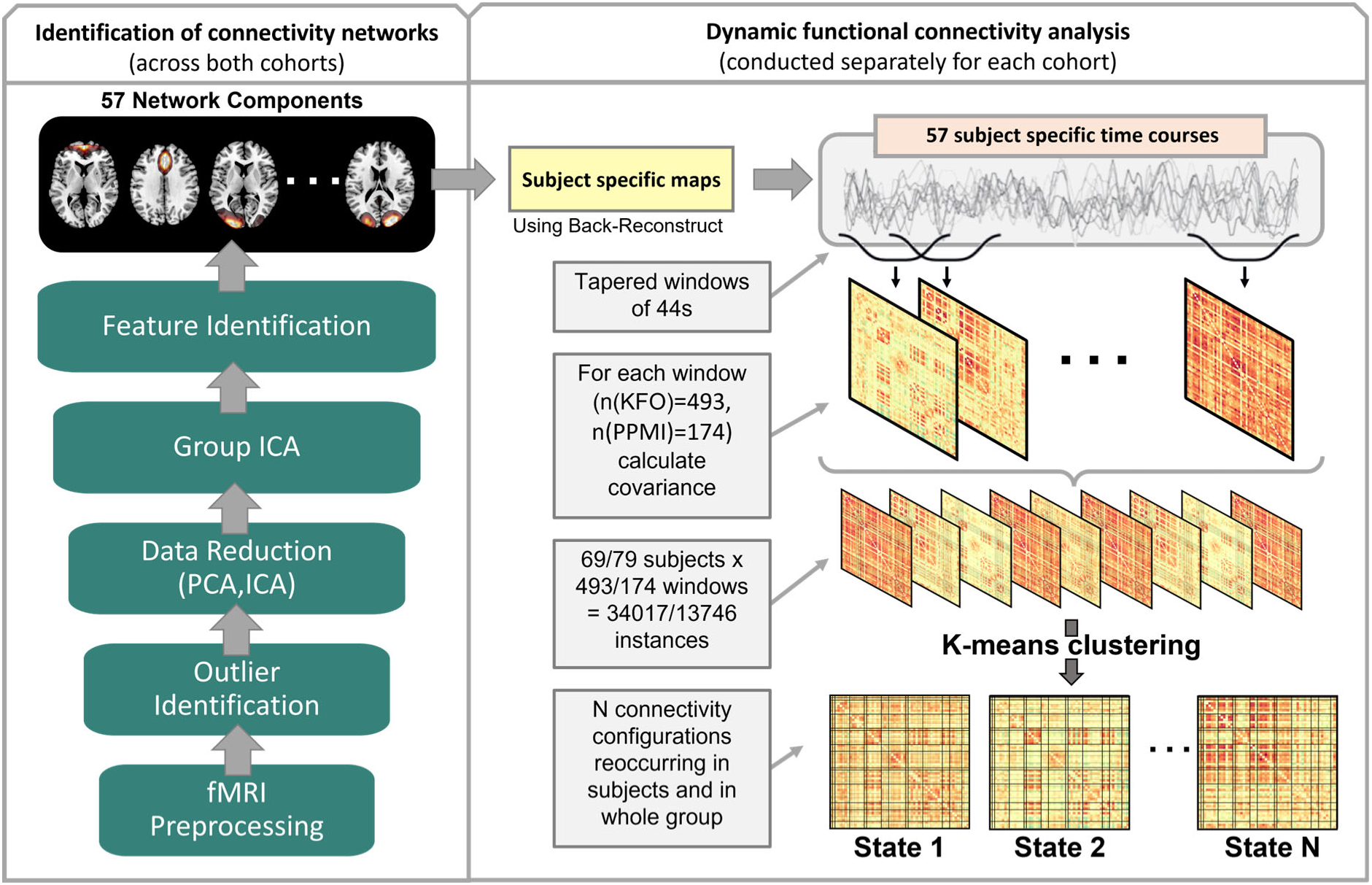
Schematic of the dynamic functional connectivity analysis pipeline: The green boxes represent the general steps to retrieve independent network components (INCs) from both cohorts. Afterward, a back reconstruction step resulted in subject-specific spatial maps and time courses for each INC. Via the sliding window approach utilizing tapered windows of 44s length, correlation matrices were obtained for each patient across the entire scan for each cohort. With the k-means clustering algorithm, the correlation matrices of the entire cohort were clustered according to similarity.

#### Sliding window approach

The sliding window approach was applied on time courses of the back-reconstructed subject-specific spatial maps obtained from the ICA, including both cohorts. Therefore, the same subject-specific maps were used in both analyses. These time courses were analyzed in frames of a defined length, called windows. The first window was set at the beginning of a time course and was then moved along the data by a predefined number of data points, defined as step variable. For each window, a covariance matrix was calculated, portraying the connectivity as covariance (correlation) between all INCs at that particular time. Each covariance matrix was composed of (57 × (57-1))/2 = 1596) features. Following previously established methods (Allen et al. 2014), a tapered window of 44s (57 TRs in KFO and 18 TRs in PPMI) was used, which was convolved with a Gaussian kernel of 3 TR. The window was moved by 1TR steps along the 617 (KFO) and 210 (PPMI) -TR scan, resulting in 493 (KFO) and 174 (PPMI) consecutive windows and their corresponding functional connectivity (FC) estimates (covariance matrices). Prior to further analysis, the FC estimates were Fisher Z transformed to improve the normality of Pearson’s r distribution. To investigate the potential effect of TR, we matched the TRs of the two samples by down-sampling the KFOs TR to 2.328 ms (average of three scans with a TR of 0.776 ms). The results were comparable for manipulated and unmanipulated TR. Hence, the TRs for the current analysis were not altered.

#### Clustering analysis

We applied the k-means clustering algorithm to all windowed FC estimates to assess reoccurring FC patterns. For each cohort, the clustering algorithm was initially performed on a defined subset of exemplary windows, chosen as the windows with local maxima in FC variance, and repeated 500 times on this set. The resulting centroids (cluster medians) were then used to initialize the clustering of all data on 69 subjects times 493 windows, equating to 35.496 instances in the case of KFO and 79 subjects times 174 windows, equating to 13,746 instances in the PPMI cohort. The similarity between each FC estimate and the cluster centers was determined using the L1 distance function (Aggarwal et al., 2001). The optimal number of clusters (k) was defined using the Elbow criterion, which independently yielded four valid clusters (*k*=4) in both cohorts. Hence, based on the similarity of each FC matrix with the four obtained cluster centroids, all respective FC matrices were then categorized into one of the given four clusters (states). Reproducibility of the estimated states has previously been validated, showing that k-means clustering results yielded reproducible cluster centroids from both analyses with bootstrap resamples and split-half samples of subjects (Allen et al., 2014).

#### Global state characteristics

To derive and compare the connectivity pattern associated with a state, connectivity matrices were calculated using in-house Python scripts (Damaraju et al., 2014): First, a subject median was computed for each state based on the subject windows assigned to that state. Second, subject medians for each state were used to derive median connectivity matrices for each state per cohort (see Fig. 3). A state’s connectivity strength was defined as the absolute sum of all correlation values in its median connectivity matrix. Notably, only correlation values higher or respectively lower than 10% of the highest and lowest correlation values were included to reduce the impact of noise on our measure. To define the proportionate similarity between two states, the Manhattan distance was calculated for the two median connectivity matrices of these states. A state’s occurrence was defined as the percentage of windows assigned to it. In an attempt to further characterize the connectivity pattern entailed in a state, we reported between-network correlations that stood out upon visual inspection.

**Figure 3:**
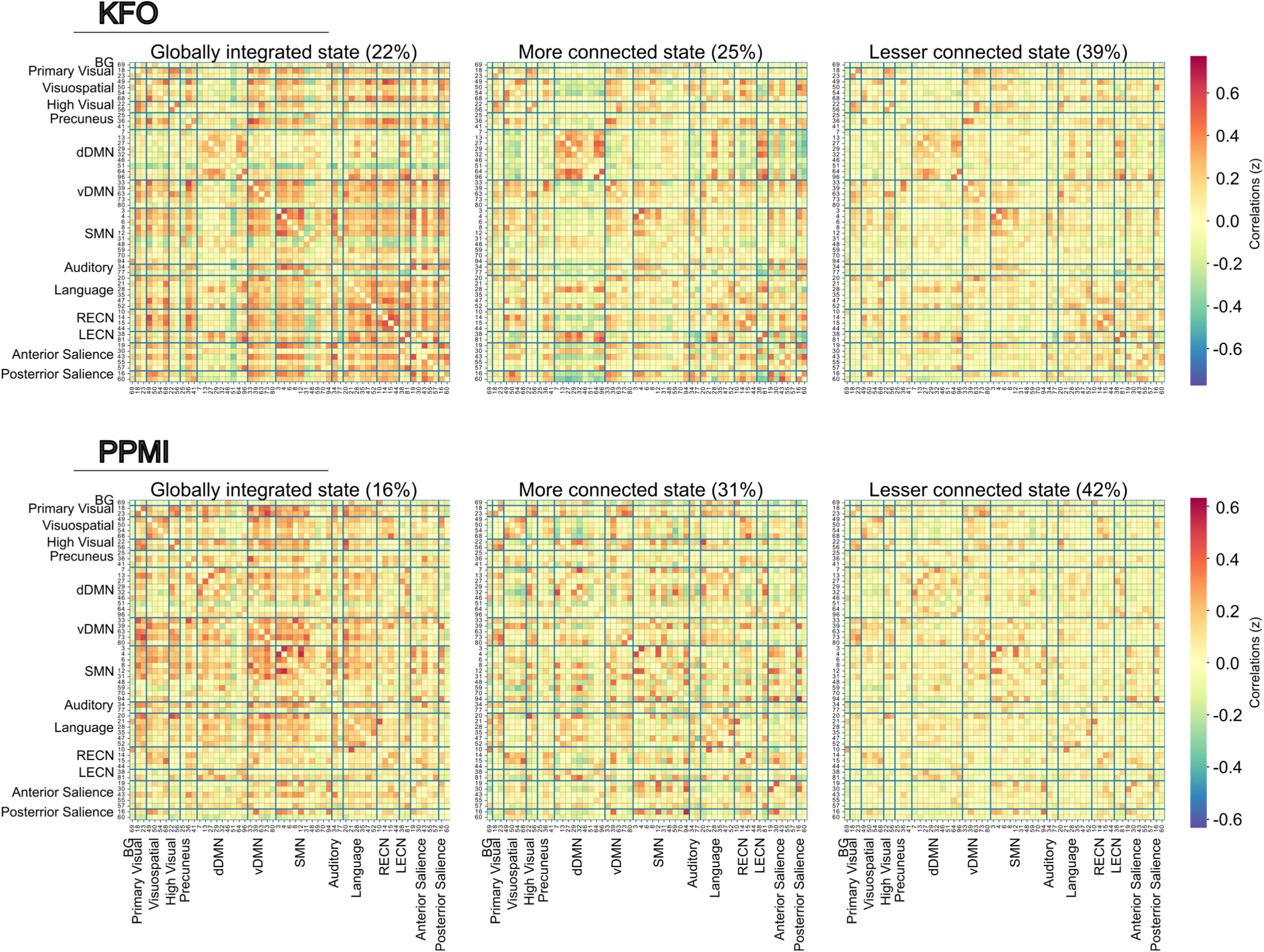
Dynamic Functional connectivity states along the whole sample (k =3) for each cohort. Each cluster is summarized in one FC map portraying FC within and between resting state networks (RSNs) (portioned by thick blue lines) in pairwise correlations. The correlations were Fisher z transformed and averaged across all subjects, then inverse z-transformed for display. The total percentage of windows assigned to the state is shown. The value of correlations is represented in the color bar; the red color represents positive and blue negative correlations. The diagonal represents the correlations of the subnetwork by itself and is thereby one.

#### Calculation of temporal properties

Temporal properties of the dFC analysis were extracted, analyzed, and plotted for each cohort separately using in-house Python scripts: Based on the categorized data obtained by the clustering analysis in GIFT, state transition vectors were created, representing the assigned state for each window for each participant. To examine the temporal properties, we assessed four variables: average dwell time, total number of transitions, bi-directional transitions, and state attendance. The average dwell time was defined as the mean time spent in a particular state. We also assessed the total number of transitions between any states and the number of bi-directional transitions between specific pairs of states. A state was considered as attended if at least one window of that participant was assigned to that particular state.

### Statistical analyses

All statistical analyses were carried out using SPSS (SPSS Statistics ver.28, IBM Corp.) and Python (Spyder; Python ver. 3.10). Given that assumptions of normality and homoscedasticity were not met in terms of the dFC variables per cohort, non-parametric tests were applied. As the PD and control groups did not differ in age and sex inside and across cohorts, the models were not corrected for these covariates. Considering states with generally low attendance are a potential cause for unreliable statistics, we only included states attended by more than 50% of the cohort’s subjects. Hence, state four (23 %) of the KFO cohort and state two (23%) of the PPMI cohort were excluded from the statistical analyses. Corrections for multiple comparisons were carried out using the Dunn and Šidák criterion (Šidák, 1967).

#### Group comparisons of dFC measures

Mann-Whitney U tests were performed to compare the PD and control groups in terms of variables derived from dFC analysis (i.e., average dwell time, fraction time, and number of transitions). For categorical variables like state attendance, Chi-squared tests were utilized.

#### Correlation analyses – dFC measures, dopamine synthesis, and behavior

To assess whether dFC measures were associated with cognitive and motor performance, MMSE and UPDRS-III scores were correlated with average dwell time and the number of total and bi-directional transitions using two-sided rank correlation tests (Kendal Tau). Besides, to examine whether temporal properties of a distinct state were associated with striatal dopamine signal, rank correlations were carried out between the mean synthesis capacity of the putamen and caudate VOIs and the dFC measurements.

## Results

### Group characteristics

In both cohorts (see Table 1), we found no significant differences between the control group and the PD group in terms of sex, handedness, age and MMSE. In both cohorts, PD subjects had significantly fewer years of education and lower putaminal dopamine synthesis capacity or dopamine transporter availability, respectively.

Across cohorts, PD subjects did not show significant differences regarding sex, handedness, age and UPDRS-III OFF scores. The PD group included in the KFO study had fewer years of education, lower MMSE scores, and a longer disease duration compared to the PD group of the PPMI cohort. Across cohorts, the KFO control group also showed lower values on the MMSE compared to PPMI controls. Sex, handedness, age, and education did not differ between the KFO control and the PPMI control group.

### Dynamic functional connectivity

#### Global state characteristics

In both cohorts, we identified three distinct connectivity states that reoccurred in individuals throughout a resting state scan and were present across subjects (see Figure 3).

For the KFO cohort, the two most frequently occurring states (a total of 64% of all windows) showed the most similar connectivity patterns (Manhattan distance of 119) across all state pairs (See Fig 3. top middle and top right pane). They mainly differed in overall connectivity strength. One state was clearly more interconnected with absolute correlation sums of 211.6 and 146.1, respectively. Hence, this state will be referred to as the more interconnected (MC) state, while the other state will be referred to as the lesser connected (LC) state of this state pair. Upon visual inspection, the pair can be characterized by strong correlations of the default mode network (DMN) with itself and parts of the left executive network (LECN) and the language network. The other state occurred less frequently (a total of 22% of all windows) and exhibited high correlations for the ventral DMN with the visual networks, the language network and parts of the SMN, the auditory network, and the salience networks (see Fig 3 top left pane). As this state showed high interconnectivity for all RSN despite the auditory network, the dorsal DMN, and parts of the SMN, we will refer to this state as the globally integrated (GI) state.

Among all states of the PPMI cohort, two states exhibited the highest similarity measured by Manhattan distance (Manhattan distance of 107) (see Fig 3. bottom middle and bottom right). These states occurred by far the most (total of 73 % of all windows) and primarily differed in connectivity strength (absolute correlation sum of 107.8 vs. 178.0). Hence, one state of the pair will be referred to as the more connected (MC) state, while the other will be termed the lesser connected (LC) state. The other state occurred least often (a total of 16% of all windows) and showed a very distinctive connectivity pattern (see Fig 3 bottom left pane). On visual assessment, this state presented relatively high connectivity between the DMN, the primary visual, and the somatomotor networks. Following the framework in the KFO data, it will be referred to as the globally integrated (GI) state.

The similarity between the sorted state profiles between both cohorts was described by Manhattan distance as follows: The highest similarity for the GI state and the LC state of the KFO cohort were the GI and the LC state of the PPMI cohort, respectively. For the MC state of the KFO, the LC state of the PPMI cohort was most similar.

#### Group differences in temporal properties

No significant group differences were found in terms of average dwell time in the KFO cohort. In the PPMI cohort, controls appeared to spend more time in the GI state than PD patients (U(N=79) = 361.5, z = −1.88 p= .033). However, this did not hold up against correction for multiple comparisons (k=3: adj. p=.017). For the remaining states, no significant differences in dwell time were found in PPMI (see Fig 4A).

**Figure 4:**
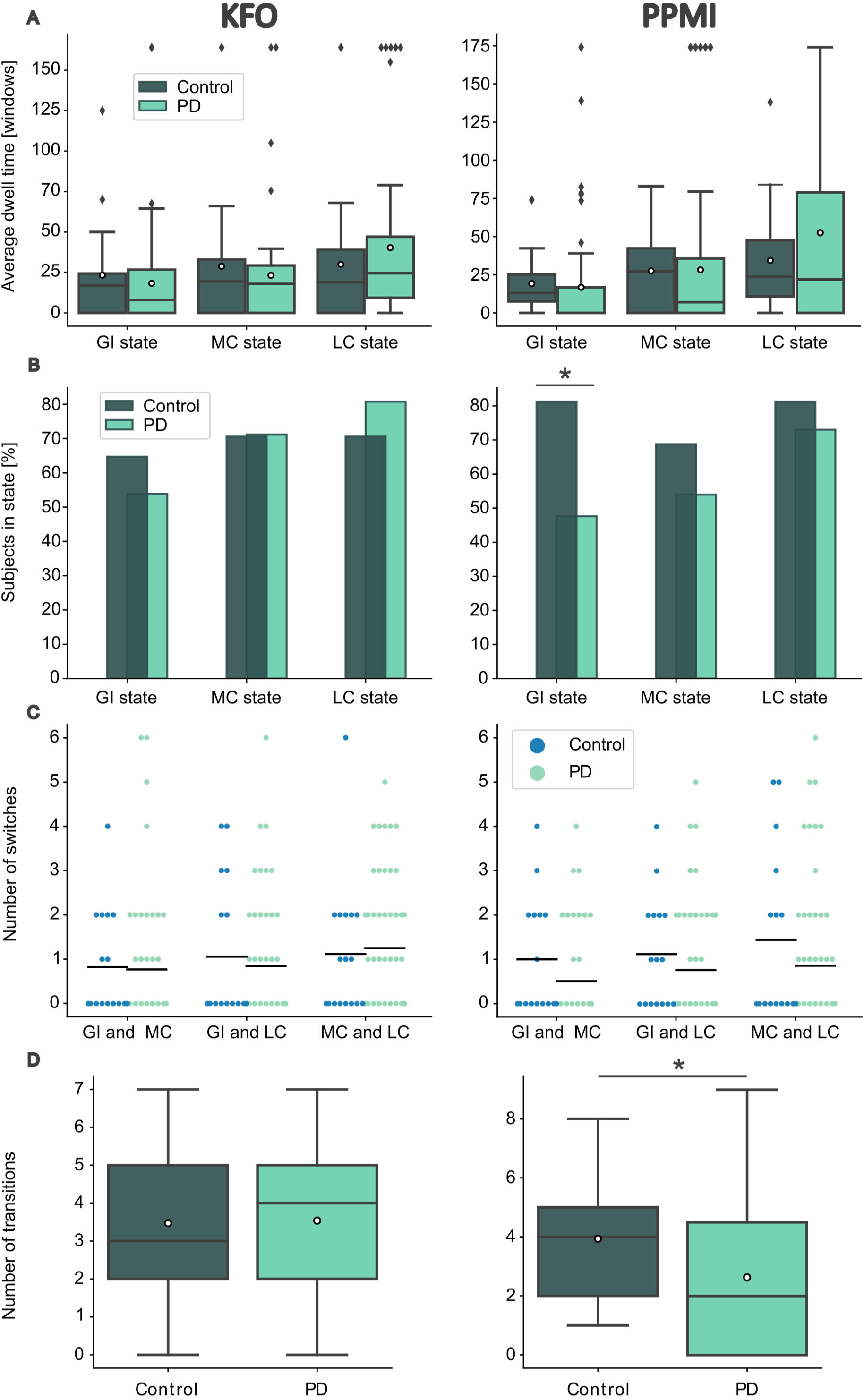
Temporal properties of dFC states for Parkinson’s disease and healthy control groups. **(A)** Boxplot portraying the distribution of the average time spent in the globally integrated (GI), the more connected (MC), and the lesser connected (LC) state for each cohort. Each colored box represents the interquartile range (IQR), with the median plotted as a thin line and the mean represented by a white dot. The whiskers specify minimum and maximum. Outliers are shown in the form of diamonds. Time is specified in windows. **(B)** Percentual proportion of subjects having dwelled in the GI, the MC, and the LC state at least once during a scan. A Chi-squared test tested for between- group differences. **(C)** Group-specific number of transitions between the pairs of states GI and MC, GI and LC, and MC and LC. The black lines indicate the mean values. Within-group differences were tested via the Friedmann test. **(D)** Boxplot portraying the distribution of the total number of transitions between all states. Mann-Whitney U-tests tested for between-group differences; *p<0.05.

We determined the number of subjects having at least once dwelled in a state during acquisition, to compare state attendance between the groups. In the KFO cohort, no group differences were observed for state attendance. For the PPMI cohort, this yielded a higher state attendance of the GI state for the control group in contrast to the PD group (X^2^(1, N=79) = 5.82, *p*= .016) (see Fig 4B).

To determine further differences between groups, we compared the total number of transitions as well as the bi-directional transitions between pairs of states. For the KFO cohort, no group differences could be reported regarding total and bi-directional transitions. When assessing the distribution of the total number of transitions in the PPMI cohort, controls exhibited a higher number of transitions than PD patients in the (U(N=79) = 337.5, z = −2.06 p= .039) (See Figure 4D). In terms of bi-directional transitions, we found trends in the PPMI cohort. Here, controls indicated a higher number of transitions between the GI and the MC state (U(N=79) = 388.5, z = −1.85 p= .064) (see Figure 4C).

#### Correlation between caudal dopamine availability, dynamic functional connectivity measures, and clinical measures

##### DFC with clinical measures

We tested for an association between MMSE and UPDRS-III scores with the number of bi- directional transitions, average dwell times, and total number of transitions. In the KFO cohort, an association between UPDRS-III scores and the number of bi-directional transitions between the GI and the LC state (τ_b_(N=52)= −.230; *p* =.036) was observed in the PD group. However, this did not hold against the Dunn and Šidák criterion (k=2: adj p=.025). For the MMSE scores, no associations with dFC were found in the KFO cohort. In the PPMI cohort, a correlation analysis revealed significant negative associations between UPDRS-III motor scores and average time spent in the GI state (τ_b_(N=63) = −.281; *p* =.003), the total number of transitions (τ_b_(N=63)= −.308; *p* = .001) and number of bi-directional transitions between the GI and the LC state (τ_b_(N=63)= −.237; *p =*.019) in the PD group. Moreover, we found two additional negative associations between UPDRS-III scores and the average time spent in the MC state (τ_b_(N=63) = −.194; *p* = .038) and the number of bi-directional transitions between the MC and the LC state (τ_b_(N=63)= −.210; *p* =.035). However, these did not survive the correction for multiple comparisons (k=2: adj. *p*= .025). In the PPMI cohort, no associations of dFC variables with MMSE were found.

##### Dopamine with clinical measures

In a subsequent step, we tested whether presynaptic dopaminergic insufficiency of the two striatal VOIs was associated with either cognitive scores or motor function in the PD groups. The correlation analysis yielded a significant negative correlation between the mean putamen values and UPDRS-III OFF scores in the KFO cohort (τ_b_ (N=41) = −.256 *p*= .048) and the PPMI cohort (τ_b_ (N=63) = −.189 *p*= .031). Both results did not withstand correction for multiple comparisons (k=2: adj. *p*= .025). In both cohorts, we did not find an association between cognitive scores of the MMSE and mean dopamine scores in the putamen or the caudate.

##### Dopamine with dFC

As the average dwell time, the number of bi-directional transitions, and the number of total transitions correlated with UPDRS-III off scores in the PPMI cohort, we tested the association between striatal dopamine synthesis capacity and the three dFC properties across the PD group in both cohorts. Inside the KFO PD group, no such associations between dFC variables and dopamine synthesis capacity could be reported. Inside the PPMI cohort, average time spent in the GI state correlated with the mean DaT availability in the putamen (τ_b_(N=63)=.213, *p*= .023) and the caudate (τ_b_(N=63)=.209, *p*= .025). The number of total transitions was also associated with mean DAT availability in the putamen (τ_b_(N=63)=.350, *p*<.001) and the caudate (τ_b_(N=63)=.251, *p*=.007). In terms of bi-directional transitions we observed an association of the number of transitions between the GI and MC state with mean DAT availability in the caudate (τ_b_(N=63)=.236, *p*=.02). We further found an association of bi- directional transitions between the GI and MC state with the mean DAT availability in the putamen (τ_b_(N=63)=.201, *p*=.042). This association, however, did not survive correction for multiple comparisons (k=2: adj. p=0.025).

## Discussion

The primary aim of this study was to elucidate dopamine’s influence on whole-brain network dynamics in PD. This study used two matched cohorts to assess the reproducibility of the dFC results. Ultimately, we found PD-specific network dynamics in the PPMI cohort: A decrease in the total number of transitions and a lower overall presence of the GI state. Since these specific features of dFC were negatively associated with UPDRS-III scores, we suspect a high transition frequency and an increased presence of a GI state to facilitate motor performance. As these have further been shown to depend on striatal dopamine availability, dopamine appears to play an essential role in mediating those large-scale network dynamics in early PD patients. Notably, our results were not stable across cohorts, suggesting that the dFC alterations depend on cohort composition.

Employing dFC on two cohorts, we identified three connectivity states in both cohorts that followed the same pattern. In both cohorts, there were two states of high total occurrence, showing the highest similarity regarding connectivity patterns of all three states: One was highly connected, while the other was less connected. The last state, less frequent in both cohorts, was characterized by a more global, integrated connectivity pattern. Hence, using two independent k-means clustering approaches, we disentangled two similar connectivity cluster profiles for both cohorts, enabling us to compare inside group differences between cohorts. In the PPMI cohort, we found the overall expression of these states and the dynamic transition between states to be group-dependent. For the KFO cohort, however, these could not be replicated. Hence, if not explicitly stated, all the findings discussed below were found only inside the PPMI cohort.

First, PD individuals of the PPMI cohort showed overall decreased transitions between states. Indeed, most of the previous studies in whole-brain dFC in PD reported differences in the total number of transitions (Cordes et al., 2018; Díez-Cirarda et al., 2018; Fiorenzato et al., 2019). However, their results are conflicting. While one study reported a higher number of transitions in PD (Díez-Cirarda et al., 2018), two other studies showed a lower transitioning frequency in PD patients compared to controls (Cordes et al., 2018; Fiorenzato et al., 2019). Interestingly, in two of these studies, this trend solely depended on cognitive status (Díez-Cirarda et al., 2018; Fiorenzato et al., 2019). Although our findings suggest that PD patients exhibit limited dynamics in dwelling behavior, these results were not associated with deteriorating cognitive abilities, as reported by Fiorenzato and colleagues (Fiorenzato et al., 2019), or overall greater cognitive flexibility, as suggested by Nomi and colleagues(Nomi et al., 2017). Against this body of studies, we found features of dFC in the PPMI cohort to be associated with motor performance. PD patients who, in total, transitioned less between states showed higher motor impairments according to the UPDRS-III. This result contradicts previous observations linking better motor performance with reduced transition behavior (Kim et al., 2017). Of particular importance and in contrast to the previously mentioned papers, dFC was assessed in patients OFF medication in our cohorts. This approach could mean that our results better represent PD’s actual decompensated state of brain networks.

Moreover, we observed the same relationship when comparing the specific transitioning patterns between groups. Individuals who specifically transitioned more often between the GI and the LC state showed better motor performance. In contrast, most of the studies mentioned above identified only two states in their study or did not investigate differences in transitioning patterns if more states were present. Hence, we are the first to report associations of specific transitioning patterns with motor performance in PD. These associations suggest that overall high network dynamics and dynamics between particular connectivity configurations are relevant in motor control. Thus, in PD, the underlying mechanisms facilitating these dynamic transitions between states of global integration and segregation seem to be impaired. Intriguingly, our results in PPMI further imply that this ability to generally transition between states and to specifically transition between globally segregated and integrated states depends on the integrity of the striatal dopaminergic system. This notion is based on two findings. First, we observed, trend significant, the well-known relationship between the decline in DaT signal and the increase in the UPDRS-III motor score (Benamer et al., 2000; Pirker, 2003; Seibyl et al., 1995) inside the PPMI PD group and the KFO cohort. Second, in PPMI PD patients, the ability to dynamically transition between all states, and the ability to transition specifically between the GI and LC state, depended on caudal and putaminal DaT availability.

Further, fewer PD patients of the PPMI group visited the globally integrated state at least once during an 8-minute acquisition compared to controls. Hence, many PD individuals missed spending time in a state characterized by high interconnectivity between networks. Together with our previous results indicating that dynamic transitions between states of global integration and segregation are impaired in PD, these results extend the notion that PD patients may have problems transitioning specifically into a state of global integration. Interestingly, another study longitudinally investigating dFC changes in PPMI data revealed a decrease in dwell time in a state similar to the GI state and an increase in dwell time for a state corresponding to the lesser connected state(Cao et al., 2023). Their results suggest that hypocoupling worsens in the course of PD (Cao et al., 2023). We only found a trend significant decrease in average dwell time in the globally integrated state for PD. Yet, our other findings in the same cohort suggest this worsening of hypocoupling to be caused by an inability to transition into a state with high interconnectivity. Central to this, in our study, the average time spent in the GI state and the number of bi-directional transitions between the GI and the LC state were positively associated with motor performance in the PPMI cohort. This suggests the increased presence of hypocoupling and decreased dynamics between integrated and segregated connectivity configurations to be related to poor motor output. Similar associations were reported in whole-brain dFC before; however, again, the opposite effect was reported (Kim et al., 2017).

Nevertheless, another recent study showed levodopa intake to increase the dwell time in a state with the strongest functional coupling between the fronto-parietal network (FPN) and the SMN (Chen et al., 2021). OFF medication, this mean dwell time was reduced again in the same PD patients (Chen et al., 2021). Notably, this dopamine-associated change in mean dwell time was negatively associated with UPDRS-III scores, highlighting that altered dFC features are associated with motor performance and, particularly dopamine-dependent. In our PPMI cohort, we could reproduce these findings with a direct measure of dopaminergic impairment, as the mean dwell time in the globally integrated state was positively associated with the mean DaT signal in the putamen and the caudate. While Chen and colleagues’ study was focused on dynamics between the SMN and networks involved in top-down motor control, our study assessed network dynamics on a whole-brain level. Hence, our results obtained in the PPMI cohort extend their view of SMN and FPN stability to be dopamine-dependent, pointing to the fact that global integration of the SMN with other non-motor associated networks is dopamine-dependent and relevant to sustaining proper motor output.

According to our results in the PPMI cohort, we suspect that in unmedicated, early PD patients, network changes are affiliated with motor function rather than cognitive performance. In particular, the overall presence of globally interconnected states and high dynamics towards a state of high interconnectivity seem to facilitate motor performance. As these have further been shown to be dependent on striatal dopamine availability, dopamine appears to play an essential role in mediating those large-scale network dynamics in early PD patients. With these findings, we provide a novel mechanism for how dopamine might modulate motor control in PD and highlight the potential of investigating large-scale network dynamics in a disease context. In particular, investigating bi-directional transitions between connectivity states, which have never been assessed in PD, provided novel insights into how dopamine might contribute to motor functions.

Importantly, although the state profiles of both cohorts were very well matched, most of the results reported above were not reproduced in the KFO cohort. In this section, we will discuss potential reasons for that. First, we will examine whether the methodological approach itself may be the root cause of our different results in both cohorts. Second, we will assess how demographical differences and cohort composition might have fueled differences in dFC results across cohorts.

Due to the unconstrained nature of resting state experiments, the concept of rsfMRI itself may appear challenging when reproducing results in two cohorts. In rsfMRI, the participants get no specific task other than keeping their eyes open. Hence, we cannot ensure that the same brain regions are comparatively engaged across participants with no task at hand. Nonetheless, static RSNs were proven to have high levels of reproducibility across imaging sessions (Biswal et al., 2010; Shehzad et al., 2009) and different subjects (Damoiseaux et al., 2006; Shehzad et al., 2009). Through our data-driven approach that included both cohorts simultaneously, we established RSNs that were similarly expressed across all individuals of both cohorts. This enabled us to compare differences between groups inside the cohort and the obtained results across cohorts. By choosing comparatively similar sample sizes for controls and PD patients in both cohorts, we ensured that individuals of one particular cohort did not dominate the RSN composition. However, the clinical presentation of PD is heterogeneous. Previous reports indicate a difference in DMN and striatal connectivity in PD patients with tremor-predominant and primarily akinetic rigid symptoms (Karunanayaka et al., 2016; Zhang et al., 2015). Based on this, although PD groups were matched across cohorts, RSN connectivity might have varied in both PD groups due to different subtype ratios.

Furthermore, inside the cohorts, group sizes deviated highly from each other. Controls comprised 25.4% of the PPMI and 32.7% of the KFO cohort. Hence, overall RSN architecture could have been driven by PD RSN composition. Despite that argument, during our definition of meaningful components, the RSNs identified by the ICA considerably overlapped with RSNs previously established in groups of younger healthy controls (Shirer et al., 2012). Refining RSNs for each group in each cohort separately would resolve these two issues. However, they would critically dampen the between-group and especially the between-cohort comparability of our dFC results.

Likewise, the dFC approach might be susceptible to technical differences in data acquisition. Hence, although the same analysis pipelines were used, alterations in data acquisition may interfere differently with dFC results. For instance, it has been suggested that the duration of a dFC-appropriate resting state acquisition should be at least 10 min (Hindriks et al., 2016). Hence, the extended acquisition time in the PPMI resting-state protocol of 8.5 minutes might have been better suited to a dFC approach than the KFO’s 8-minute acquisitions. Moreover, the cohort’s acquisition paradigms varied drastically in TR. Hence, the analysis was carried out for the KFO cohort separately, once with a TR adjusted to the PPMI dataset and once with an unadjusted TR. Although these two approaches equally did not produce any sign group differences regarding dFC variables for the KFO cohort, we cannot rule out that the different TRs might affect dFC results. Nevertheless, as changing the KFO’s TR by downsampling would entail a significant manipulation of the input data, we kept the data as they were. Interestingly, a preceding study provided evidence of the reproducibility of basic connectivity patterns amidst inter-regional connections over 7500 scans composing “probably” mixed independent resting-state datasets of healthy and diseased individuals (Abrol et al., 2017). However, all these scans followed the same acquisition protocol, and only 8 of these scans deviated in TR from the other scans. Vital to mention at this point is a recent study investigating the relationship of dFC alterations with non-motor symptoms of PD in a PPMI cohort. This study used the same analysis pipeline with slightly altered settings (50s windows, Yeo atlas). Intriguingly, this study found astonishingly similar state patterns to those we achieved for the PPMI cohort in our approach (Cao et al., 2023). This demonstrated the reproducibility of state patterns across different analysis parameters and cohort composition in one of our cohorts. Further, this high similarity in sorted state patterns signifies that joining both cohorts in one ICA did not alter the state pattern composition of our PPMI cohort.

Further, differences in cohort composition could be why the results observed in the PPMI cohort were not reproduced in the KFO cohort. Comparing the demographics of both PD groups revealed two things: First, PPMI included de-novo PD individuals, while KFO included early PD patients. Hence, PD individuals of the KFO cohort were more advanced in the progress of the disease than the ones of the PPMI cohort. Second, KFO PD individuals showed lower levels of education and lower scores in cognitive tests despite inherently difficult comparisons between countries. According to preceding studies, PD-specific alterations of dFC increase with time (Cao et al., 2023) and cognitive impairment (Díez-Cirarda et al., 2018; Fiorenzato et al., 2019). Hence, we expected KFO PD individuals to be even more likely to show between-group differences than PPMI individuals. Further, as previously elaborated and as intended, dopamine medication exerts a normalizing effect on functional connectivity strength in static FC (Berman et al., 2016; Tahmasian et al., 2015) and dFC (Chen et al., 2021). Although in both cohorts, dopaminergic medication was discontinued before the fMRI scans, the long-term effects of dopaminergic medication cannot entirely be excluded (Cilia et al., 2020). Significantly, the medication discontinuation paradigms varied between cohorts. While in KFO, conservative periods of a minimum of 12h discontinuation for dopamine replacement and a maximum of 78h for extended-release agonists were established, in PPMI, more moderate ranges of 6h to 28h (average 16h) of medication discontinuation were present. Hence, both cohorts might have imbalanced discontinuation periods, entailing differently pronounced long-term effects of medication on rsfMRI results.

Considering all this, we can summarize that we identified differences in cohort composition and acquisition parameters that could have caused variation in the between-group differences observed in both cohorts. Yet, we did not identify one specific feature in methodology or cohort composition that we consider highly influential. We suspect these differences in the results to be driven by differences in cohort composition that were not assessed within the framework of this study, like disease subtype ratio or lifestyle factors. This again underlines the need for deep phenotyping in neuroimaging datasets, as a more detailed description of the cohorts and more significant group sizes would have been highly relevant to performing an effective group matching.

## Conclusion

Our findings in the PPMI cohort illustrate that PD patients exhibit altered network dynamics compared to healthy controls. These alterations are defined by a lower prevalence for integrated connectivity states and decreased transition frequency between a globally integrated and a lesser connected state. With these findings, we supply a potential mechanism for how dopaminergic neurodegeneration drives global changes in connectivity underlying motor dysfunction in PD. However, we could not reproduce these whole-brain dFC results in another cohort. Therefore, it will be valuable to reassess these findings and their replicability in two bigger, effectively matched cohorts. Furthermore, it will be interesting to investigate whether the reported impairment of network dynamics is already observable in prodromal stages of PD, such as in patients with REM-sleep behavioral disorder.

## Acknowledgments

We are grateful for the patience and commitment of every participant participating in the study.

## Funding

This work was supported by the Deutsche Forschungsgemeinschaft (DFG, German Research Foundation) – Project-ID 431549029 – SFB 1451.

## Data availability

The data that support the findings of this study are available on request from the corresponding author.

## Financial Disclosures

AA reports no conflicts of interest. MH reports receiving research funding from the German Research Foundation (DFG). T. van Eimeren received honoraria, stipends or speaker fees from the Lundbeck Foundation, Gain Therapeutics, Orion Pharma, Lundbeck Pharma, Atheneum, and the International Movement Disorders Society. He receives materials from Life Molecular Imaging and Lilly Pharma. He owns stocks of the corporations NVIDIA, Microsoft and I.B.M. GRF serves as an editorial board member of Cortex, Neurological Research and Practice, NeuroImage: Clinical, Zeitschrift fuer Neuropsychologie, DGNeurologie, and Info Neurologie & Psychiatrie; receives royalties from the publication of the books Funktionelle MRT in Psychiatrie und Neurologie, Neurologische Differentialdiagnose, and SOP Neurologie; receives royalties from the publication of the neuropsychological tests KAS and Koepps; received honoraria for speaking engagements from Deutsche Gesellschaft fuer Neurologie (DGN) and Forum fuer medizinische Fortbildung FomF GmbH.

## Authors’ Roles

AA, MH, and TVE were responsible for the design and conduct of the analysis. AA wrote the first draft of the manuscript, which co-authors carefully reviewed.

